# Leptin signaling regulates physiological damage and host-pathogen cooperation

**DOI:** 10.1101/2020.08.24.264648

**Authors:** Karina K. Sanchez, Katia Troha, Sarah Stengel, Janelle S. Ayres

## Abstract

To combat infections, hosts employ a combination of antagonistic and cooperative defense strategies. The former refers to pathogen killing mediated by resistance mechanisms, while the latter refers to physiological defense mechanisms that promote host health during infection independent of pathogen killing, leading to an apparent cooperation between the host and the pathogen. Previous work has shown that leptin, a pleiotropic hormone that plays a central role in regulating appetite and energy metabolism, is indispensable for resistance mechanisms, while a role for leptin signaling in cooperative host-pathogen interactions remains unknown. Using a mouse model of *Yersinia pseudotuberculosis* (*Yptb*) infection, the causative agent of Far East scarlet-like fever, we unexpectedly found that genetic inhibition of leptin signaling conferred protection from *Yptb* infection due to increased host-pathogen cooperation rather than greater resistance defenses. The protection against *Yptb* infection was not due to differences in food consumption, lipolysis or fat mass. Furthermore, we found that the survival advantage was associated with increased liver damage and dysfunction. Our work reveals an additional level of complexity for the role of leptin in infection defense and suggests that in some contexts, in addition to tolerating the pathogen, tolerating organ damage and dysfunction is more beneficial for survival than preventing the damage.

## INTRODUCTION

Leptin is an adipose tissue hormone that acts via its receptor (LEP-R) in the brain to regulate energy balance and neuroendocrine function (Pan and Myers, 2018). Circulating leptin levels communicate the state of body energy repletion to the central nervous system in order to suppress food intake and promote energy expenditure (Deck et al., 2017; Pan and Myers, 2018). When leptin is present at normal levels, it supports energy utilization on a variety of essential processes, including those that support the immune system. Conversely, leptin deficiency results in increased appetite and altered energy expenditure—leading to obesity—as well as decreased immune function (Deck et al., 2017; Francisco et al., 2018; Pan and Myers, 2018). Indeed, leptin deficiency has historically been associated with increased susceptibility to a variety of infections (Hsu et al., 2007; Mancuso et al., 2002; Radigan et al., 2014; Tschop et al., 2010).

To combat infections, hosts use a combination of evolved antagonistic and cooperative defense strategies (Ayres, 2020; Raberg et al., 2009; Schneider and Ayres, 2008). The former refers to pathogen killing mediated by resistance mechanisms, while the latter refers to physiological defense mechanisms that foster host health during infection independent of pathogen killing, leading to an apparent cooperation between the host and pathogen (Ayres, 2016; Ayres, 2020). The traditional view of leptin’s role in infection defense is that leptin is required to mediate resistance mechanisms and pathogen killing. For example, phagocytosis, an intricate immune process through which cells ingest and eliminate pathogens, is severely impaired in the absence of leptin (Dayakar et al., 2016; Moore et al., 2003; Tschop et al., 2010). In a mouse model of *Streptococcus pneumoniae* infection, leptin-deficient mice exhibited defective alveolar macrophage phagocytosis. This was associated with reduced bacterial clearance in the lungs and increased lethality. Administration of exogenous leptin to leptin-deficient mice during the infection improved phagocytosis, bacterial clearance, and animal survival (Hsu et al., 2007). Leptin also modulates the function of several other innate immune cells, thus influencing the innate immune response against pathogens. For instance, leptin promotes chemotaxis and the release of pathogen-killing reactive oxygen species in neutrophils (Alti et al., 2018; Souza-Almeida et al., 2018). Similarly, it supports the development and activation of natural killer cells (Tian et al., 2002). Additionally, leptin exerts a variety of effects on the adaptive immune system, such as increasing the expression of adhesion molecules by CD4^+^ T cells and promoting a bias toward TH1-cell responses in memory CD4^+^ T cells (Fernandez-Riejos et al., 2010). While leptin has been shown to play a large role in the promotion of resistance mechanisms, and therefore antagonistic defenses, the role of leptin in cooperative defense strategies remains unexplored.

Cooperative defense mechanisms include disease tolerance mechanisms, which act to prevent/alleviate physiological damage during infection and enable the host to endure such damage should it occur, and anti-virulence mechanisms, which dampen microbial and host derived virulent signals and behaviors that contribute to disease pathogenesis (Ayres, 2016; Ayres, 2020; Troha and Ayres, 2020). A general theme that has emerged from recent studies is that physiological defense mechanisms largely involve metabolic processes induced during infection (Luan et al., 2019; Rao et al., 2017; Sanchez et al., 2018; Schieber et al., 2015; Wang et al., 2016). For example, infection-induced tissue iron sequestration and insulin resistance promoted host-pathogen cooperation in an enteric pathogen model (Sanchez et al., 2018). Other studies have found that sickness-induced anorexia can either promote or hinder cooperative defenses during infection (Ayres and Schneider, 2009; Rao et al., 2017; Wang et al., 2016). In another study, a microbiome *E. coli* strain promoted disease tolerance during enteric and pulmonary infections by protecting mice from infection-induced wasting (Schieber et al., 2015). Given the metabolic nature of the physiological defense mechanisms identified thus far and the role that leptin plays in the maintenance of systemic metabolic homeostasis, we reasoned that leptin may play a major role in regulating physiological defenses and host-pathogen cooperation.

While disease tolerance and anti-virulence defenses are evolved mechanisms to promote physiological defenses and host-pathogen cooperation, a loss of antagonistic defenses can also yield an apparent cooperation and protection from damage during infection. This occurs when the cost of mounting the resistance response is greater than the benefit of killing the pathogen (Ayres, 2020). Such costs come in the form of immunopathology that occurs as a consequence to the resistance response. Typically, the loss of such responses will provide protection from tissue/organ/physiological damage and cooperation, leading to a survival advantage despite elevated levels of the pathogen. We have less of an appreciation of the inherent costs to mounting physiological defense responses and whether there are contexts for which enduring tissue/organ/physiological damage is more beneficial than the costs of mounting a disease tolerance or anti-virulence response.

*Yersinia pseudotuberculosis* (*Yptb*) is a Gram-negative, extracellular bacterium that is pathogenic to humans and mice (Davis, 2018; Galindo et al., 2011). *Yptb* is transmitted via the fecal-oral route. Upon oral infection, *Yersinia* traverses the M cells of the intestine. Once across, it infects the underlying lymphoid tissues (Peyer’s patches [PP] and mesenteric lymph nodes). Infection typically causes self-limiting gastroenteritis and mesenteric lymphadenitis. In mice, and occasionally in humans, *Yptb* spreads to the liver and spleen and causes systemic disease leading to sepsis (Davis, 2018; Galindo et al., 2011; Mikula et al., 2012). The ability of *Yptb* to multiply in lymphoid tissues and spread to systemic organs depends on the presence of pYV, a virulence plasmid that encodes type III secretion system (T3SS) and several effector proteins called Yersinia outer proteins (Yops). The Yops function to inhibit phagocytosis, downregulate proinflammatory cytokines, and induce apoptosis in target cells, among other functions (Cornelis et al., 1998; Mohammadi and Isberg, 2009; Viboud and Bliska, 2005; Wang et al., 2014b). Given that *Yersinia* infection is known to increase transcription of the leptin receptor gene (*Lepr*) in mice and that several studies have established a strong link between *Yersinia* virulence and core metabolism (Fischer et al., 2019; Heroven and Dersch, 2014), *Yptb* is an excellent pathogen to elucidate the relationship between leptin, metabolism, and cooperative defenses.

Here, we describe a role for leptin signaling in the regulation of physiological defenses and host-pathogen cooperation. In agreement with previous reports (Hsu et al., 2007; Mancuso et al., 2002), we found that *Lep^ob^* and *Lep^db^* mice that are deficient for leptin signaling had impaired resistance defenses against *Yptb* and reduced inflammation. Unexpectedly, however, we found that despite the increased pathogen burdens, leptin signaling deficient mice had a survival advantage compared to wild type animals, indicating leptin signaling regulates host-pathogen cooperation. Furthermore, we found that the survival advantage was associated with increased liver damage and dysfunction. Our work reveals an additional level of complexity for the role of leptin in infection defense and suggests that in some contexts, in addition to cooperating with the pathogen, tolerating organ damage and dysfunction is more beneficial for survival than preventing the damage.

## RESULTS

### Leptin deficiency promotes health and survival of *Yptb* infection

We began by characterizing the clinical course of disease for *Yptb* infection in C57Bl/6 mice to understand how this infection influences metabolic health of the host. Oral infection with this enteropathogen resulted in 100% mortality by 5 days post-infection (**Figure 1A**). This mortality was associated with substantial weight loss and infection-induced anorexia. By 72 hours post-infection, *Yptb*-infected mice lost ~10% of their body weight (**Figure 1B**). The reduction in weight may partly be explained by the presence of infection-induced anorexia in these mice. Anorexia is a sickness behavior commonly induced by infection. It is characterized by a reduced motivation to consume food and triggers a fasted-like state in the host (Hart, 1988). Over the first 72 hours post-infection, *Yptb*-infected mice progressively lowered their food intake, with day 3 consumption equaling, on average 25% of their daily food intake under uninfected conditions (**Figure 1C**). In sum, our results indicate that oral infection with *Yptb* in WT mice leads to metabolic pathologies in the host including anorexia and rapid weight loss followed by death.

**Figure 1.**
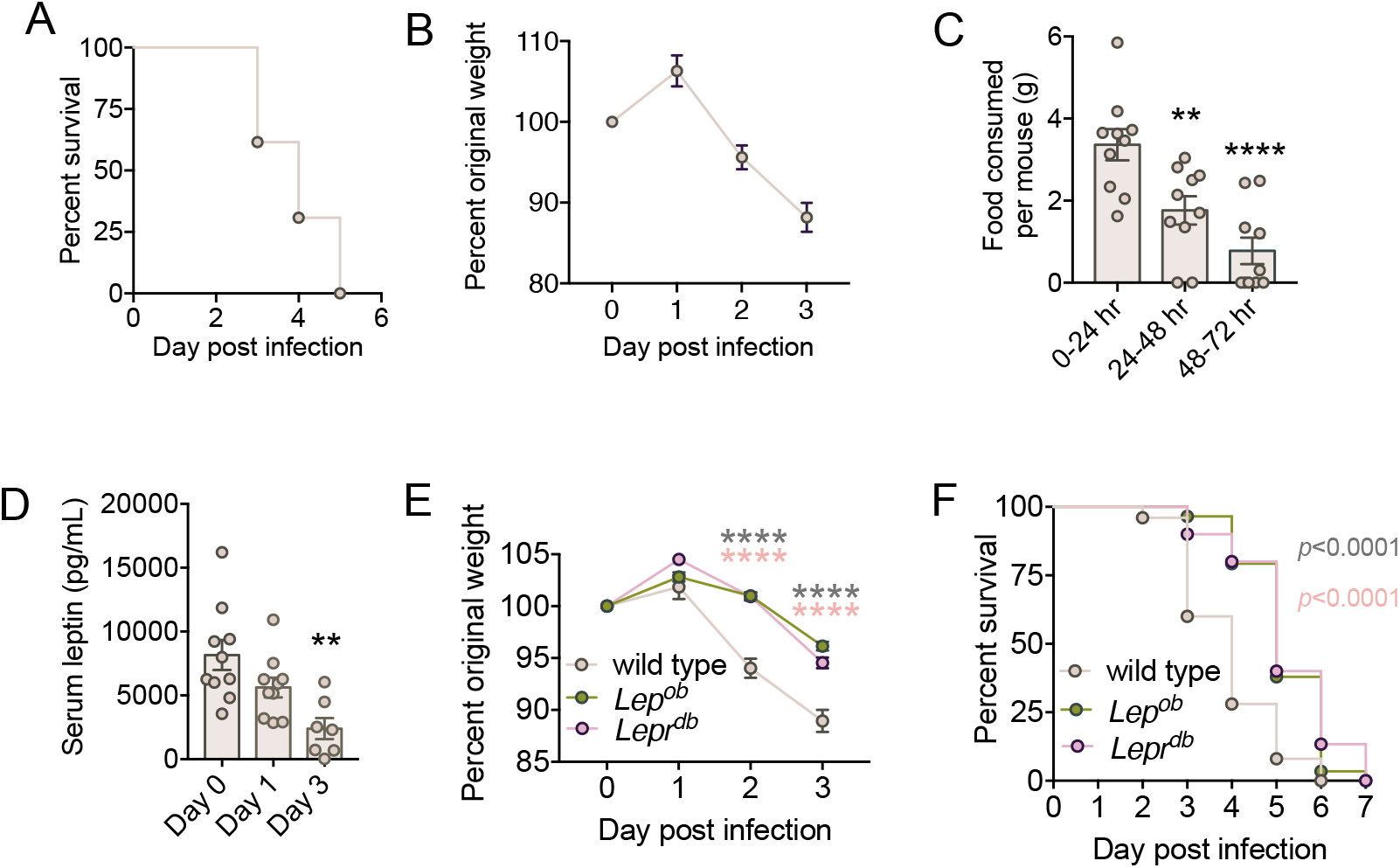
Loss of leptin signaling promotes health and survival during *Yptb* infection. (A) Survival of WT mice infected with *Yptb*. n = 13 mice. (B) Weight loss of WT mice following *Yptb* infection. n = 10 mice, at day 0 and 1, n = 9 at day 2 and n = 8 at day 3 due to death from infection. (C) Food intake of WT mice infected with *Yptb*. n = 10 mice per group. Significance noted indicates that timepoint compared to 0-24hr. ** p = 0.0087, ****p<0.0001. One way ANOVA with post Tukey test. (D) ELISA measurements of serum leptin in WT mice. n = 7-10 mice per group. **p=0.0016 compared to day 0. One way ANOVA with post Tukey test. (E) Weight loss of WT, *Lep^ob^*, and *Lepr^db^* mice after *Yptb* infection. n = 25-30 mice per group. ****p<0.0001 in One way ANOVA with post Tukey test. (F) Survival of WT, *Lep^ob^*, and *Lepr^db^* mice following *Yptb* infection. n = 25-30 mice per group. ****p<0.0001 in a Log-rank test. Error bars represent +/- SEM.

Leptin is an appetite-suppressant hormone whose levels are known to increase systemically in animal models of lipopolysaccharide (LPS)-induced anorexia (Grunfeld et al., 1996; Mastronardi et al., 2001). To begin to understand the relationship between leptin and the presence of infection-induced anorexia in our *Yptb* infection model, we quantified serum leptin in *Yptb*-infected mice. Surprisingly, we found that circulating levels of leptin declined over the course *Yptb* infection in C57Bl/6. The decline was apparent as early as 24 hrs post-infection and continued to decline through 72 hours post-infection (**Figure 1D**). These data indicate that in the context of *Yptb* infection, the infection-induced anorexic response is associated with a decline, rather than an increase, in circulating leptin. The discrepancy between our results and previous work using LPS (Grunfeld et al., 1996; Mastronardi et al., 2001) may be explained by the fact that LPS administration does not fully reflect the complexity of infection with a live pathogen.

We considered the hypothesis that leptin downregulation may reflect a host defense strategy against infections, and that reduced leptin may be beneficial for the outcome of infection with *Yptb*. Therefore, we set out to determine whether the absence of leptin signaling changes the outcome of *Yptb* infection. Leptin-deficient (*Lep^ob^*) and leptin receptor-deficient (*Lepr^db^*) mice develop obesity due to hyperphagia that is sustained by a defective leptin circuitry (Wang et al., 2014a). When we infected *Lep^ob^* and *Lepr^db^* mice with *Yptb*, we found that compared to wild type infected mice, these mutant mice were protected against the weight loss induced by *Yptb* infection. While infected wildtype mice lost ~10% of their body weight by 72 hrs post-infection, infected *Lep^ob^* and *Lepr^db^* mice lost ≤ 5% of their body weight at this same time point (**Figure 1E**). In addition to a protection from infection-induced wasting, we found that mice deficient for leptin signaling had a significant survival advantage compared to infected wild type mice. The median time to death for *Lep^ob^* and *Lep^db^* mice was extended by 25% (**Figure 1F**). Altogether, our data demonstrate that leptin signaling deficiency protects from wasting and death during *Yptb* infection and suggest that lowering systemic leptin may be an adaptive strategy that promotes host defense during infection.

### Increased feeding does not promote survival of *Yptb* infection

Infection-induced anorexia is associated with distinctive changes in the expression of feeding and neuronal activation genes—termed the anorexic program—in the hypothalamus, a region of the brain that plays a central role in the regulation of food intake and energy balance (Rao et al., 2017). To confirm that the reduced food consumption exhibited by *Yptb* infected wild type mice was in fact due to anorexia, we determined whether *Yptb* infection induces the molecular anorexic program in mice. We quantified the mRNA expression of the feeding-associated genes *Lepr, Npy, Fos, Socs3, Pomc*, and *Agrp* in the hypothalamus using qRT-PCR. Compared to uninfected controls, we found that the mRNA expression of *Agrp, Socs3, Fos*, and *LepR* significantly increased, while the mRNA expression of *Npy* and *Pomc* significantly decreased in *Yptb*-infected mice (**Figure 2A**). In accordance with previously published results (Rao et al., 2017; Schwartz et al., 1997; Sucajtys-Szulc et al., 2010), these hypothalamic transcriptional changes are in line with the presence of anorexia, indicating that *Yptb*-infected mice not only display anorexic feeding behavior but also possess molecular markers characteristic of anorexia.

**Figure 2.**
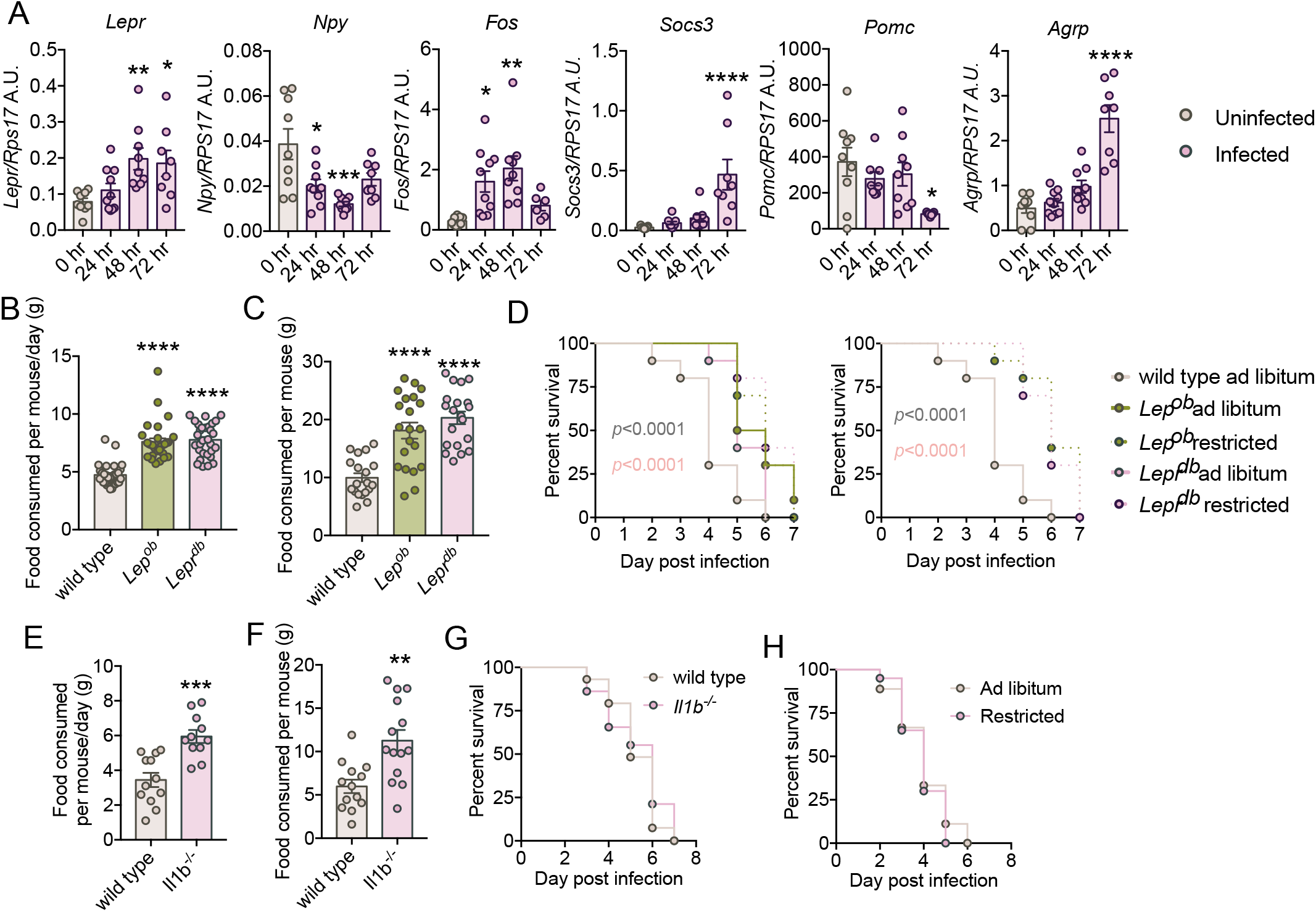
Increased feeding does not improve survival of *Yptb* infection. (A) Quantification of feeding-associated genes *Lepr, Npy, Fos, Socs3, Pomc*, and *Agrp* in the hypothalamus of *Yptb*-infected WT mice using qRT-PCR. n = 6-10 mice per group. *p<0.05 **p<0.01****p<0.0001 vs uninfected in a One way ANOVA with post Tukey test. (B) Average daily food intake of WT, *Lep^ob^*, and *Lepr^db^* when uninfected. n = 29-30 mice per group. ****p<0.0001 vs uninfected in a One way ANOVA with post Tukey test. (C) Total food consumption per mouse over the first 3 days of infection. Data for WT, *Lep^ob^*, and *Lepr^db^* are shown. n = 19-21 mice per group. ****p<0.0001 vs uninfected in a One wat ANOVA with post Tukey test test. (D) Survival of infected WT, *Lep^ob^*, and *Lepr^db^* mice under *ad libitum* and food-restricted conditions. n = 10 mice per group. Log rank analysis. The curves for WT *ad libitum* and restricted mutant conditions are the same for left and right panels. Shown on left without mutant *ad libitum* conditions for easier comparison. (E) Average daily food intake of WT and *Il1β*^-/-^ mice uninfected. n = 11-12 mice per group. ***p=0.0002 in an unpaired t-test. (F) Total food consumption per mouse over the first 3 days of infection for WT and *Il1β^-/-^* mice during *Yptb* infection. n = 13-14 mice per group. **p=0.0015 in an unpaired t-test. (G)Survival of WT and *Il1β^-/-^* mice infected with *Yptb*. n = 29 mice per group. (H) Survival of *Yptb*-infected WT mice under *ad libitum* and food-restricted conditions. n = 18-20 mice per group. Error bars represent +/- SEM

We previously demonstrated that infection with the enteropathogen *Salmonella enterica* serovar Typhimurium induces the anorexic program in the hypothalamus in mice resulting in sickness-induced anorexic behavior and reduced food consumption. We demonstrated that this behavioral response triggered increased pathogen virulence during *S*. Typhimurium infection (Rao et al., 2017). Considering that *Yptb* infection also induces an anorexic signature and resulting anorexic response in mice (**Figure 1C**) and that *Lep^ob^* and *Lepr^db^* mice exhibit hyperphagic behavior when uninfected (Wang et al., 2014a) and (**Figure 2B**), we hypothesized *Lep^ob^* and *Lepr^db^* consumed more food than WT mice during a *Yptb* infection. Consistent with our hypothesis, we found that *Lep^ob^* and *Lepr^db^* mice consumed twice the amount of food (~20 grams v. ~10 grams) as WT mice over the first three days of the infection (**Figure 2C**). Taken together, these results indicate that the characteristic hyperphagia of *Lep^ob^* and *Lepr^db^* mice is extended to infected conditions.

We next tested our hypothesis that the sustained hyperphagia of *Lep^ob^* and *Lepr^db^* during *Yptb* infection was necessary to confer protection against the infection in mice deficient for leptin signaling. To do this, we performed pair-wise feeding studies where we infected WT, *Lep^ob^*, and *Lepr^db^* mice with *Yptb* and restricted the diet of *Lep^ob^*, and *Lepr^db^* each day post infection so that they were allowed to consume comparable amounts of food as WT mice each day post-infection. While food-restricted *Lep^ob^* and *Lepr^db^* mice displayed increased survival of infection compared to *ad libitum* WT mice, we did not observe any differences in survival between *ad libitum* and food-restricted *Lep^ob^* and *Lepr^db^* mice (**Figure 2D**). In sum, our data suggest that while *Lep^ob^* and *Lepr^db^* mice are hyperphagic during infection, their increased feeding does not account for the survival advantage of these mice when infected with *Yptb*.

To further corroborate that increased feeding does not lead to increased survival in the context of *Yptb* infection, we employed two alternate models. The cytokine IL-1β is a key mediator of feeding under homeostatic conditions (**Figure 2E**) and infection-induced anorexia (Dantzer, 2009). We infected WT and *Il1β^-/-^* knock-out (KO) mice with *Yptb* and measured food consumption during infection. Infected *Il1β^-/-^* KO mice exhibited a significantly higher food intake compared to infected WT mice (**Figure 2F**). These results indicate that IL-1*β* contributes to the anorexic response induced by *Yptb* infection. Next, we compared survival of infection between WT and *Il1β^-/-^* KO mice. We found that there is no survival difference between infected WT and *Il1β^-/-^* KO mice (**Figure 2G**). Thus, our second model (*Il1β^-/-^* KO mice) confirmed that increased feeding does not promote survival during *Yptb* infection. As a third and last approach, we hypothesized that if increased feeding serves to protect mice against *Yptb* infection, then food-restricting WT mice during infection will lower their survival. To test this hypothesis, we compared survival of infection between *ad libitum* and food-restricted WT mice. Our data showed no difference in survival between these two groups of mice (**Figure 2H**). In sum, using three distinct approaches, we demonstrated that food consumption does not change the final outcome of *Yptb* infection and that the protection mediated by Leptin signaling deficiency is not due to the hyperphagic behavior of these mice during a *Yptb* infection.

### Increased lipolysis or fat mass do not protect against *Yptb* infection

Infections cause dramatic rearrangements to the physiology of host energy stores including wasting of adipose tissue. In an MRI time course analysis, we found that *Yptb* infection in WT mice caused a rapid and robust adipose tissue wasting phenotype beginning 24 hrs post-infection (**Figure 3A**). By 72 hrs post-infection, *Yptb* infected WT mice lost 33.5% of total body fat and direct measurements of inguinal and gonadal white adipose tissue weights (IWAT and GWAT), revealed a >50% reduction in mass (**Figures 3B** and **3C**). *Lep^ob^* and *Lepr^db^* mice exhibit a higher rate of net lipolysis under homeostatic conditions compared to wild type mice (Turner et al., 2007). We hypothesized that this increased lipolysis may confer a survival advantage when animals are challenged with *Yptb*. To test whether lipolysis was necessary to protect against *Yptb* infection, we employed tissue specific knock out mice, *Pnpla2 fabp4 cre*, in which *Pnpla2*, the gene encoding adipose triglyceride lipase (ATGL) which mediates the first and rate limiting step of lipolysis is knocked out in adipocytes (Trites and Clugston, 2019). We reasoned that if increased lipolysis provided a survival advantage to *Lep^ob^* and *Lepr^db^* mice during *Yptb* infection, then lipolysis-impaired mice, such as ATGL-deficient mice, would be more susceptible to the infection. Mice deficient for ATGL function in adipocytes did not exhibit adipose tissue wasting when infected with *Yptb*, demonstrating that adipose tissue wasting during *Yptb* infection is dependent on ATGL function (**Figure 3D-F**). Our survival analysis showed comparable death kinetics for *Yptb* infected *cre*- and *cre+* mice, indicating that adipose tissue lipolysis is not necessary to alter the outcome of a *Yptb* infection (**Figure 3G**).

**Figure 3.**
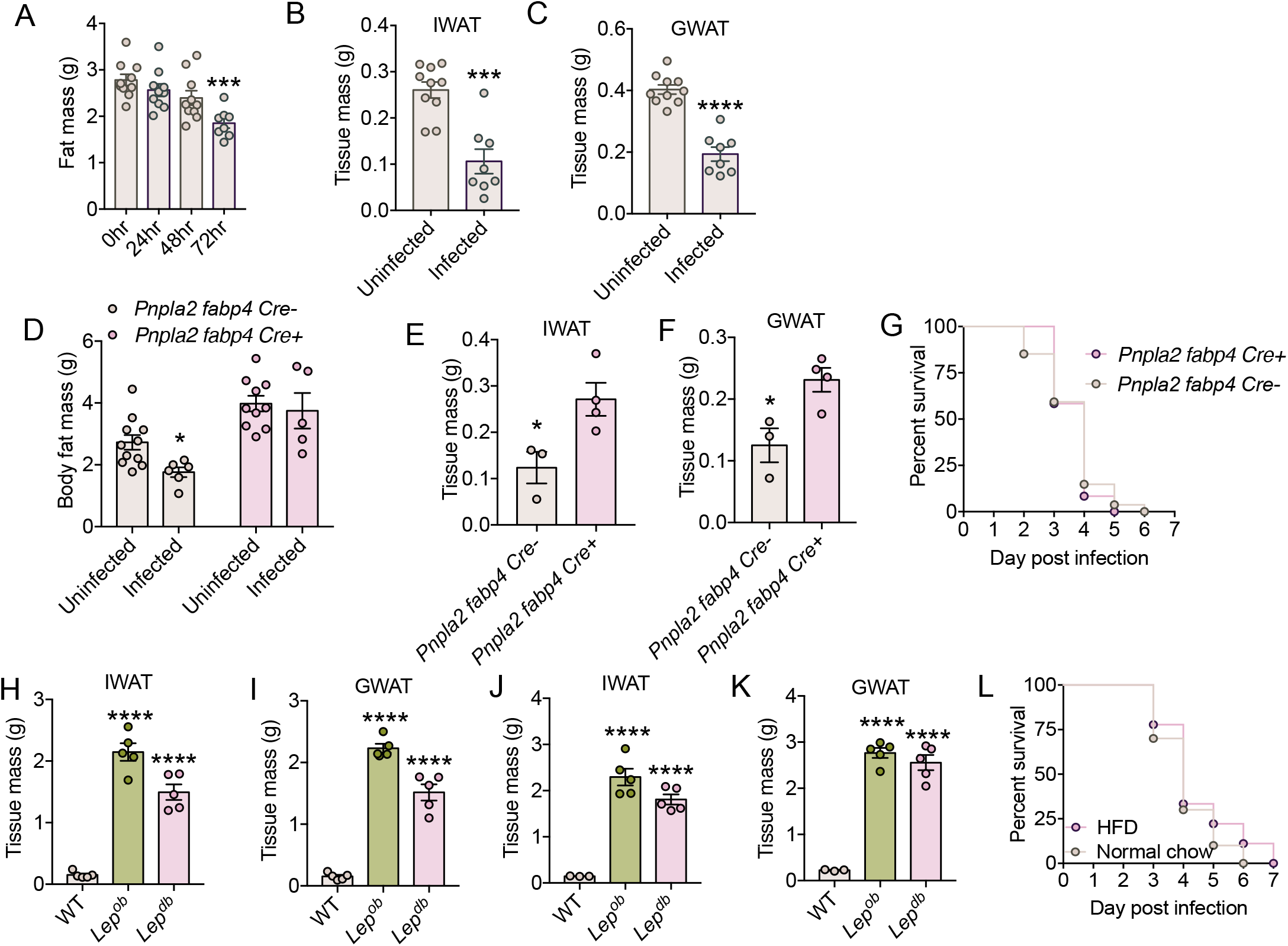
Increased fat mass and lipolysis do not confer protection against *Yptb* infection. (A-B) WT mice were infected with *Yptb* and fat mass was measured. (A) MRI analysis measuring total body fat mass; (B) IWAT and (C) GWAT were harvested from uninfected and infected mice and whole tissue weights were measured. n = 8-10 mice per group. (D-G) *Pnpla2 fabp4 cre*- and *cre+* littermates were infected with *Yptb* and fat mass and survival were measured. (D) MRI analysis of uninfected and infected mice measuring total body fat mass. n=5-11 mice per condition; (E) IWAT and (F) GWAT were harvested from uninfected and infected mice and whole tissue weights were measured. Mice that were weight matched at the time of infection were used, n=3-4 mice per condition. (G) Survival, n=12-27 mice per condition. (H-K) WT, *Lep^ob^* and *Lep^db^* mice were infected with *Yptb* and fat mass was measured. (H) IWAT and (I) GWAT were harvested from uninfected mice and tissue weights were measured, n=5 per group. (J) IWAT and (K) GWAT were harvested from infected mice and tissue weights were measured, n=3-5 mice per group. (L) WT mice fed control or HFD were infected with *Yptb* and survival was monitored, n=9-10 mice per condition. One way ANOVA with post Tukey test, t-test or Log rank analysis. Error bars +/- SEM. *p<0.05, **p<0.005, ***p<0.0005, ****p<0.0001.

*Lep^ob^* and *Lepr^db^* mice are phenotypically obese under homeostatic conditions, showing increased body fat despite having the higher rate of net lipolysis compared to wild type mice (Turner et al., 2007) and (**Figure 3H-I**). When infected with *Yptb, Lep^ob^* and *Lepr^db^* mice still have significantly higher body fat compared to infected WT mice (**Figure 3J-K**). We hypothesized that excess body fat would be sufficient to protect mice against *Yptb* infection. We compared the death kinetics of WT mice fed normal chow and WT mice raised on high fat diet (HFD) and found no differences in survival between infected normal chow mice and infected HFD mice, indicating that excess body fat is not sufficient to protect against infection (**Figure 3L**). In sum, our data demonstrate that neither increased fat mass nor lipolysis change the outcome of *Yptb* infection.

### Leptin deficiency impairs gut and GALT resistance mechanisms

Resistance defenses contribute to host health and survival of infection by mediating microbial killing mechanisms to control pathogen burdens. Previous work has shown that leptin is required for resistance mechanisms against infection. Indeed, for some infections, leptin deficiency is associated with an impaired immune response and pathogen clearance, as well as decreased survival (Hsu et al., 2007; Mancuso et al., 2002; Radigan et al., 2014; Tschop et al., 2010). We evaluated the contribution of leptin signaling to host resistance defenses against *Yptb* by measuring pathogen burdens and extraintestinal dissemination over the course of the infection. We infected WT, *Lep^ob^*, and *Lepr^db^* mice with *Yptb* and measured colony forming units (CFU) in various target organs at 24, 48, and 72 hours post-infection. CFU measurements in the small intestine (SI), cecum, and colon revealed substantially higher bacterial burdens in *Lep^ob^* and *Lepr^db^* mice compared to WT mice by 48 hours post-infection (**Figures 4A-4C**). These data suggest that resistance defenses in the intestinal tract are transiently compromised during *Yptb* infection in mice deficient for leptin signaling.

**Figure 4.**
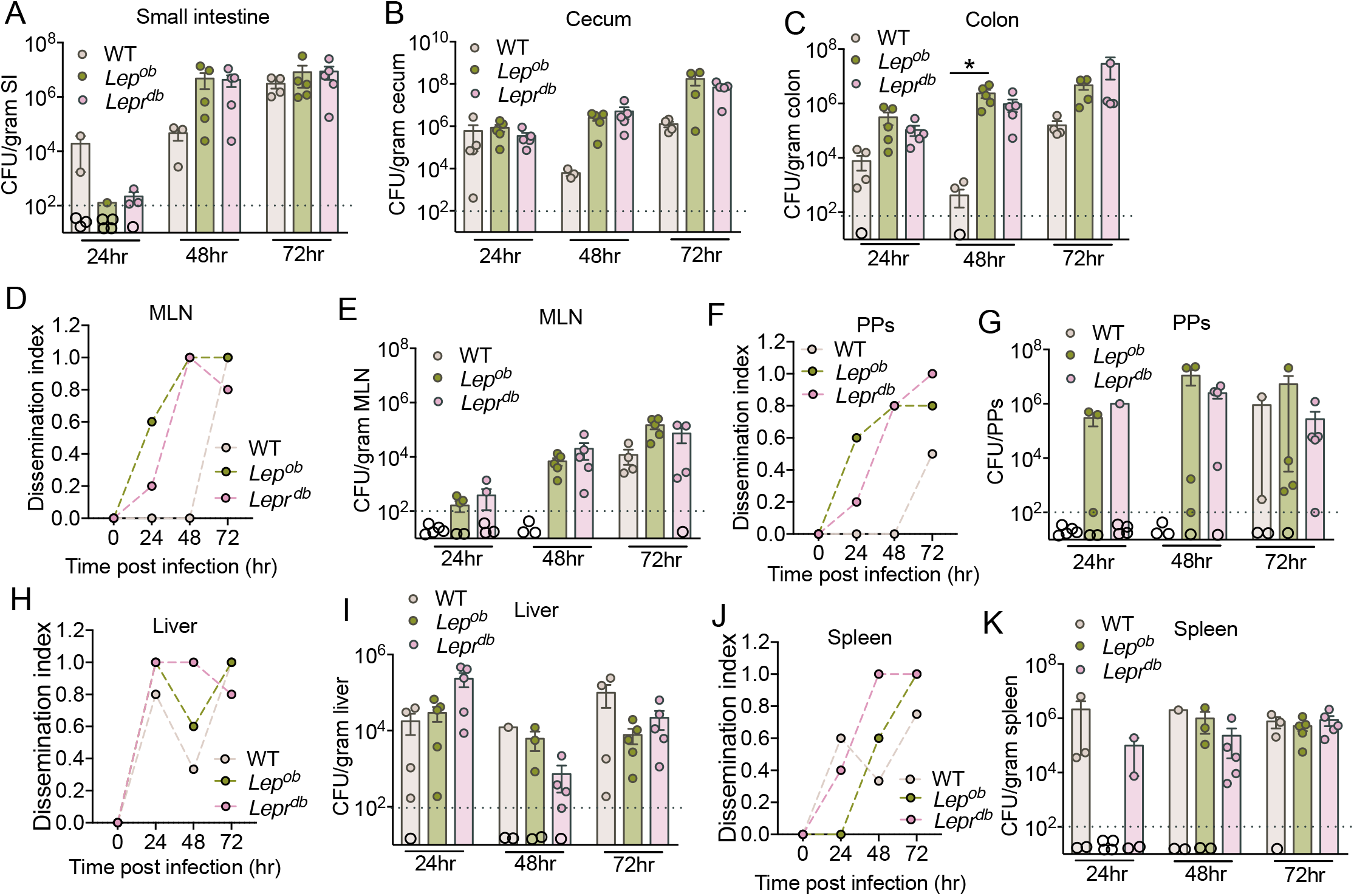
Loss of leptin impairs gut resistance mechanisms. (A-C) *Yptb* colony forming units (CFU)/gram found in the (A) small intestine; (B) cecum and (C) colon at 24, 48, and 72 hours post-infection in WT, *Lep^ob^*, and *Lepr^db^* mice. (D) Mesenteric lymph node (MLN) dissemination index at 24, 48 and 72 hrs post-infection in WT, *Lep^ob^*, and *Lepr^db^* mice. (E) *Yptb* colony forming units (CFU)/gram found in the MLNs at 24, 48, and 72 hours post-infection in WT, *Lep^ob^*, and *Lepr^db^* mice. (F) Peyer’s Patches (PPs) dissemination index at 24, 48 and 72 hrs postinfection in WT, *Lep^ob^*, and *Lepr^db^* mice. (G) *Yptb* colony forming units (CFU)/gram found in the PPs at 24, 48, and 72 hours post-infection in WT, *Lep^ob^*, and *Lepr^db^* mice. (H) Liver dissemination index at 24, 48 and 72 hrs post-infection in WT, *Lep^ob^*, and *Lepr^db^* mice. (I) *Yptb* colony forming units (CFU)/gram found in the liver at 24, 48, and 72 hours post-infection in WT, *Lep^ob^*, and *Lepr^db^* mice. (J) Spleen dissemination index at 24, 48 and 72 hrs post-infection in WT, *Lep^ob^*, and *Lepr^db^* mice. (K) *Yptb* colony forming units (CFU)/gram found in the spleen at 24, 48, and 72 hours post-infection in WT, *Lep^ob^*, and *Lepr^db^* mice. n=3-5 mice per group per time point. One way ANOVA with post Tukey test. Error bars +/- SEM, *p=0.02. Dotted horizontal line in (A-C, E, G, I and K) represent the limit of detection for the assay. Any mice with CFU below limit of detection are shown with an open circle below this line.

Pathogenic *Yersinia* have a pronounced tropism for lymphatic tissues, such as Peyer’s patches (PPs) and mesenteric lymph nodes (MLNs). We found more dissemination to these tissues, and in agreement with our gastrointestinal data, *Yptb* burdens in the PPs and MLNs of *Lep^ob^* and *Lepr^db^* mice were substantially higher compared to WT mice (**Figures 4D-G**). These data indicate that *Lep^ob^* and *Lepr^db^* mice have impaired resistance defenses in the gut-associated lymphoid tissues (GALT). Finally, in contrast to our gut and GALT results, we found no differences in pathogen dissemination to or pathogen burdens in the liver and spleen of *Lep^ob^, Lepr^db^* or WT mice (**Figures 4H-K**), suggesting that systemic resistance defenses are not compromised in *Lep^ob^* and *Lepr^db^* mice during *Yptb* infection. In summary, our data are in agreement with previous findings demonstrating that leptin is required for resistance defenses (Hsu et al., 2007; Mancuso et al., 2002; Radigan et al., 2014). However, we find that, in the case of *Yptb* infection, leptin signaling-deficient mice only have a gut and GALT specific resistance defects, as we did not observe differences in burdens at systemic sites in leptin deficient and WT animals. Furthermore, our data demonstrate that the survival advantage conferred by leptin deficiency is not due to a heightened resistance response to the infection.

### Loss of leptin signaling increases hepatic damage and dysfunction, but dampens inflammation

We found that the health and survival advantage in *Lep^ob^* and *Lep^db^* mice was associated with increased pathogen burdens in the gut and GALT tissues, and comparable burdens in the liver and spleen as found in WT mice infected with *Yptb*. Our data indicate that the absence of leptin signaling facilitates cooperation between the host and *Yptb*. While resistance mechanisms are important for host defense by killing the pathogen, if the costs of mounting such responses are too great, hosts that are able to cooperate with the pathogen will have a health and survival advantage compared to those that are more resistant (Ayres, 2020). A significant cost of resistance responses is immunopathology. Compromised resistance defenses are often associated with reduced inflammatory responses, which can contribute to a survival advantage by limiting immunopathology, despite elevated pathogen burdens. We hypothesized that the reduced resistance in the gut and GALT of *Lep^ob^* and *Lep^db^* would be associated with a reduced inflammatory response. We examined the levels of three pro-inflammatory cytokines that can contribute to immunopathology: TNFα, IL-1β and IL-6, and found that mice deficient for leptin signaling had dampened inflammatory responses in the MLNs compared to WT infected mice (**Figure 5A-C**), but no differences in the levels of these pro-inflammatory cytokines in any other gut tissues analyzed (**Supplemental Figure 1**). While we did not find differences in the pathogen burdens of *Lep^ob^* and *Lep^db^* mice compared to WT mice, we also measured the levels of pro-inflammatory cytokines in these tissues because leptin has pro-inflammatory properties. Consistent with this, we found significantly lower levels of TNFα and IL-6 in the liver and no differences in the levels of the cytokines measured in the spleen in WT and leptin deficient mice (**Figure 5D-F** and **Supplemental Figure 1**). These data indicate that the health and survival advantage conferred by deficiency in leptin signaling is associated with a dampened inflammatory response in the GALT and the liver.

**Figure 5.**
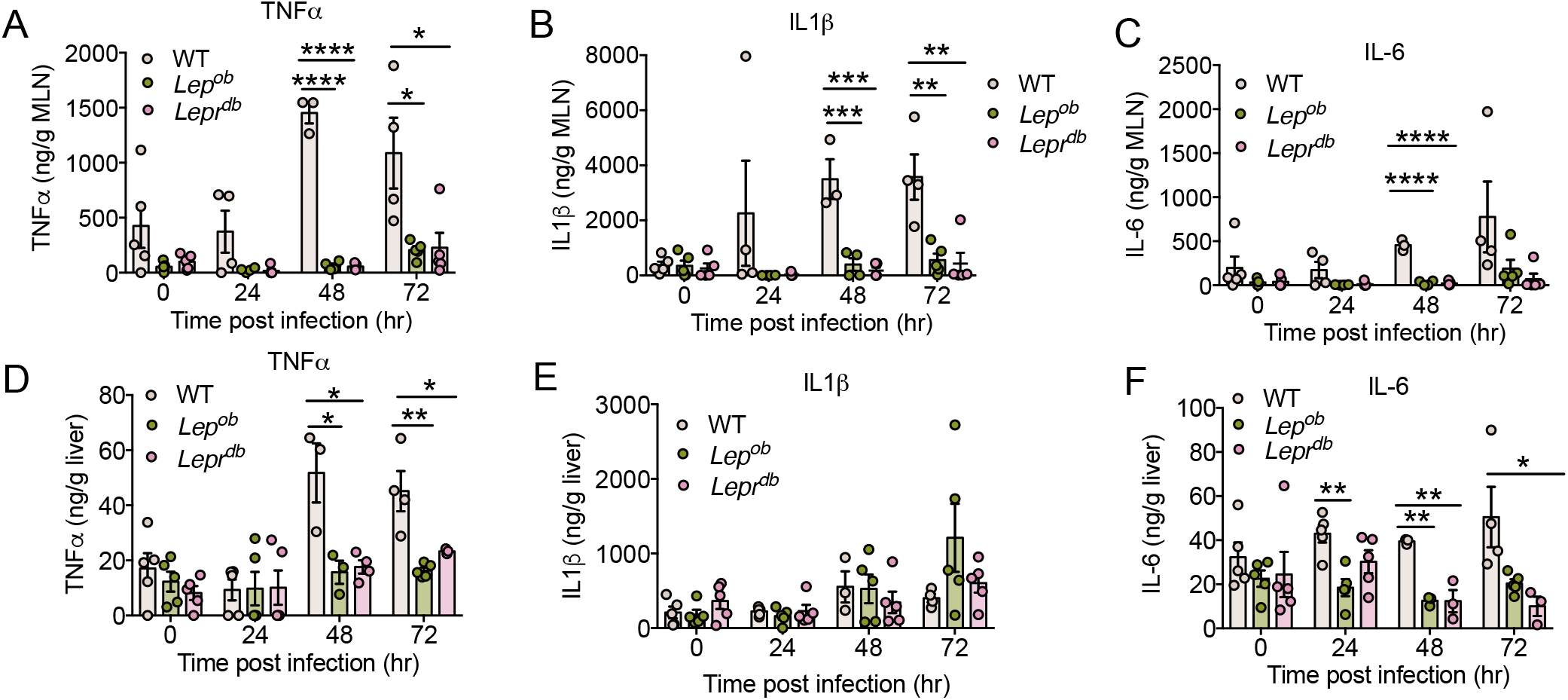
Leptin deficiency dampens inflammation during infection. WT, *Lep^ob^*, and *Lepr^db^* mice were infected with *Yptb* and proinflammatory cytokines in the (A-C) MLNs and (D-F) liver were measured by ELISA during the first 72 hrs post-infection. (A) TNFα levels in the MLNs; (B) IL-1β levels in the MLN and (C) IL-6 levels in the MLNs. (D) TNFα levels in the liver (E) IL-1β levels in the liver and (F) IL-6 levels in the liver. n = 3-5 mice per group per time point. *p<0.05, **p<0.005, ***p<0.005 and ****p<0.0001 in a One way ANOVA with post Tukey test, Error bars +/- SEM.

We hypothesized that the dampened inflammatory response and survival advantage observed in *Yptb*-infected mice deficient for leptin signaling would be associated with reduced tissue/organ/physiological damage. To test our hypothesis, we took a clinical pathology approach. *Yersinia* infection has been reported to cause both liver and kidney damage (Buhles et al., 1981; Dubois et al., 1982; Farrer et al., 1988; Kim et al., 2019; Prasad and Patel, 2018). Therefore, we measured the amount of aspartate aminotransferase (AST), alanine aminotransferase (ALT), and blood urea nitrogen (BUN) in the serum of both uninfected and infected mice. AST and ALT are markers of liver damage, while BUN is a maker of kidney function. Our data revealed that the circulating levels of both AST and ALT were significantly elevated in *Yptb*-infected mice as compared to uninfected mice (**Figures 6A** and **B**). By contrast, BUN levels were comparable between uninfected and infected WT mice (**Figure 6C**). We found that *Lep^ob^* and *Lep^db^* had significantly elevated levels of AST and ALT during infection with a modest increase in BUN levels suggesting that the absence of leptin signaling leads to increased liver damage during *Yptb* infection (**Figure 6A-C**). Furthermore, we found that creatinine and BUN:creatinine ratio decreased while bilirubin levels increased in *Lep^ob^* and *Lep^db^* infected with *Yptb* (**Figure 6D-F**), indicating reduced liver function in mice deficient for leptin signaling compared to infected WT mice. Thus, in contrast to our hypothesis, the absence of leptin signaling renders mice more susceptible to liver damage and dysfunction despite a dampened inflammatory response and increased survival of infection.

**Figure 6.**
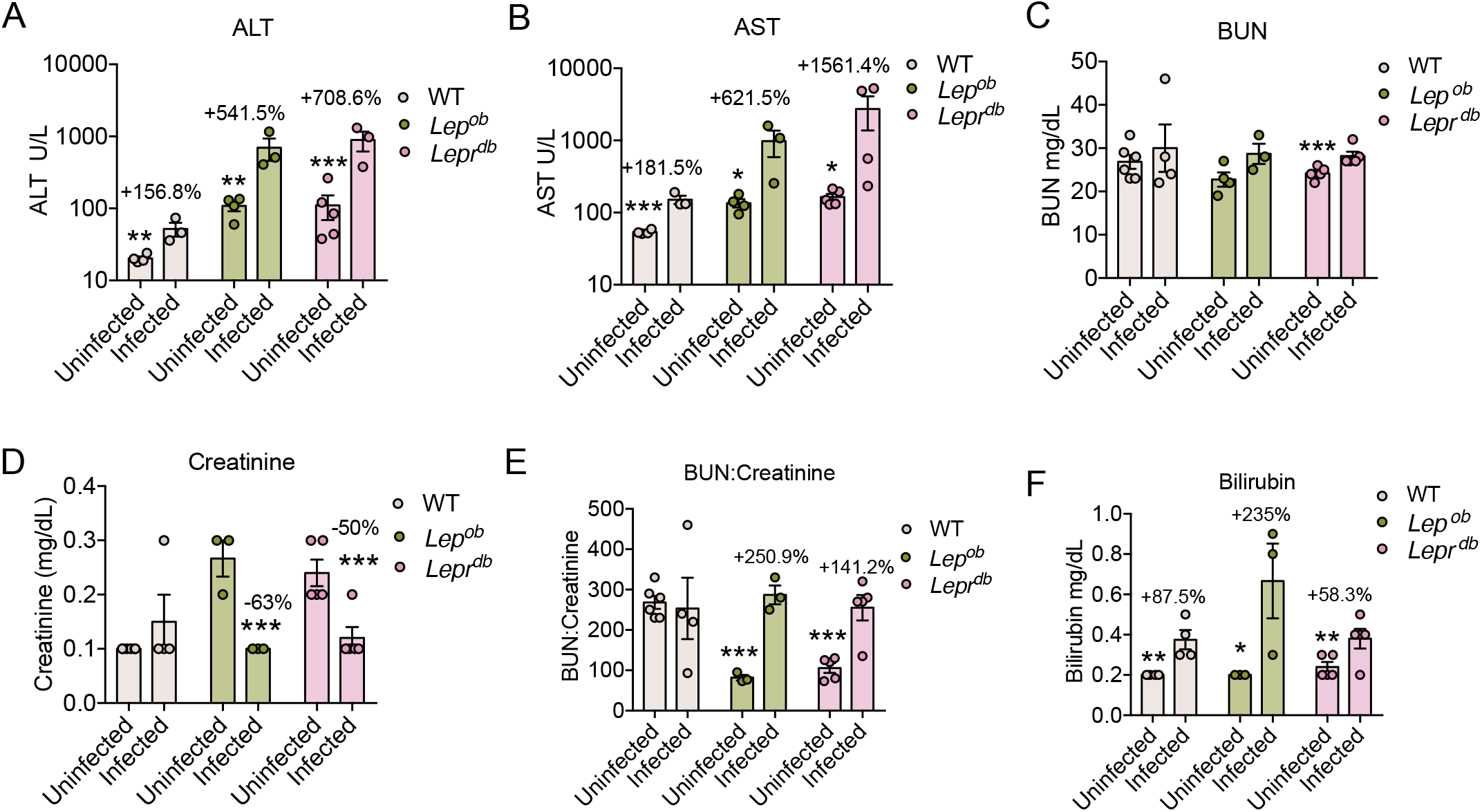
Loss of leptin signaling promotes disease tolerance during *Yptb* infection. WT, *Lep^ob^*, and *Lepr^db^* mice were infected with *Yptb* and serum levels of (A) ALT; (B) AST; (C) BUN; (D) Creatinine; (E) BUN:Creatinine ratio and (F) Bilirubin; were measured. N= 3-6 mice per condition. Unpaired t-test of uninfected vs infected for each genotype. *p<0.07, **p<0.05, ***, p<0.005 and ****p<0.0001, Error bars +/- SEM.

## DISCUSSION

The objective of this study was to investigate the role of leptin in host-pathogen cooperation (Ayres, 2016; Ayres, 2020). By taking a holistic approach, we revealed greater complexities for the role of leptin in infection defenses than previously appreciated. We found that for a single infection type: 1) the contributions of leptin signaling to host resistance defense are organ dependent; 2) the contributions of leptin signaling to the pro-inflammatory response are organ dependent and show both correlation and no correlation with pathogen burdens depending on the organ; and 3) leptin signaling protects from organ damage and dysfunction. While leptin has been long appreciated for its role in promoting pathogen clearance (Hsu et al., 2007; Mancuso et al., 2002; Radigan et al., 2014), the current study expands our understanding of leptin in host defense to include a role in the regulation of physiological defenses and host-pathogen cooperation in addition to antagonistic defenses.

We found that during *Yptb* infection, *Lep^ob^* and *Lep^db^* mice had a compromised resistance response to infection in the gut and GALT, with a dampened inflammatory response, and increased hepatic damage and dysfunction. Despite this, we found that the deficiencies in leptin signaling conferred a survival advantage for *Yptb* infection. A loss of resistance and dampened inflammation can confer a survival advantage to an infected host if the cost of mounting such responses is too costly for host health (Ayres, 2020). Typically, these costs come in the form of immunopathology, and a loss of these responses promote survival by protecting the host from tissue/organ/physiological damage. Our finding that the survival advantage conferred by a loss of leptin signaling was associated with increased hepatic damage and dysfunction was unexpected. There are several non-exclusive models to mechanistically explain our results. First, while we did not observe differences in pathogen burdens in the liver, we did find leptin deficient mice to have greater burdens of *Yptb* in the gut and GALT. The increased hepatic damage can be an indirect consequence of the decreased resistance defenses in these tissues. Second, we found reduced levels of pro-inflammatory cytokines in the liver that may be necessary for protection from hepatic damage. A third model is that leptin signaling is important for mediating protection against hepatic damage and dysfunction by a pathogen-independent and inflammation-independent mechanism. In all of these cases, the fact that we found animals that had survival advantage to have increased hepatic damage and dysfunction indicates that there are costs to protecting from this damage and dysfunction and in the context of this infection, allowing the damage to occur is less costly than protecting the host from it. While we have a good understanding of the costs associated with resistance and inflammatory responses, we do not have a good understanding of the costs for mounting tissue protective responses. It is tempting to speculate that our results may have revealed a damage allocation model in which the body allocates damage to parts of the body that are able to withstand damage to protect other parts that may be less able to withstand damage. Allocation is a common theme in infection biology in the context of resource allocation (Ayres and Schneider, 2012) and perhaps damage allocation will emerge as an important theme in infection biology with future studies.

The health and survival advantage in leptin signaling deficient animals suggests that they are able to endure the consequences of hepatic damage and dysfunction that occurs during *Yptb* infection. We suggest that this is likely due to a disease tolerance that operates to allow the host to remain healthy by enduring the damage that has occurred. We do not have a good understanding of how disease tolerance mechanisms work to facilitate health and function despite damage and dysfunction, but one likely possibility is that physiological compensation is occurring (Ayres, 2020). This involves the adaptation of other systems in the body to the new “state” to facilitate functioning and sustain health despite the damage. Future studies that are aimed at understanding how *Lep^ob^* and *Lep^db^* mice succumb to a *Yptb* infection and how this compares to *Yptb* infected WT animals will be important for understanding how deficiencies in leptin signaling influence disease course and outcome.

In our study, we revealed mechanistic insights for a number of metabolic aspects of *Yptb* infection. We found, that similar to other enteric infections (Rao et al., 2017), *Yptb* induces infection-induced anorexia and this is dependent on IL-1β. In a mouse model of *S*. Typhimurium infection, myeloid cells of the lamina propria released IL-1β that signaled to the vagus nerve to induce the anorexic response (Rao et al., 2017). Future work is needed to determine the cellular source of IL-1β that is necessary for driving the anorexic response during a *Yptb* infection and whether it is sensed peripherally or centrally to trigger the response. In the case of *S*. Typhimurium, the anorexic response led to a more severe infection course. By contrast, using dietary and genetic approaches, we found that abolishing the anorexic response had no effect on the outcome of a *Yptb* infection in mice. This is consistent with previous studies demonstrating that the effects of the anorexic response on infection outcome are infection specific (Ayres and Schneider, 2009; Wang et al., 2016). We also found that *Yptb* induces wasting of white adipose tissue over the course of the infection and we demonstrated that ATGL in the WAT is necessary to mediate lipolysis and WAT wasting during *Yptb* infection. Similar to the anorexic response, we found that inhibition of this lipolytic response had no influence on the outcome of infection. Whether adipose tissue wasting serves a role in defense against bacterial infections is not understood.

It is interesting to note that given the significant number of preceding studies to evaluate the role of leptin in infection defense, these studies, unlike ours, did not find that leptin deficiency promotes survival of pathogenic infection and host-pathogen cooperation. On the contrary, most found that the absence of leptin severely decreases survival (Hsu et al., 2007; Mancuso et al., 2002; Radigan et al., 2014; Tschop et al., 2010). We speculate that there are at least two main reasons for this occurrence. First, the evolutionary solutions that have arisen from different host-pathogen co-adaptations will have unique ways in which each pathogen interacts with its host. Another layer of intricacy adding to whether decreasing leptin is beneficial or detrimental to the outcome of specific infections comes from leptin’s dual role in inflammation. While leptin is generally viewed as a pro-inflammatory factor, acting to enhance the production of various inflammatory cytokines (Otero et al., 2006), leptin has also been reported to reduce inflammation in some infection models. For example, during *S. aureus* infection, which reduces circulating leptin in the host, administration of exogenous leptin lowers the levels of IL-6 and reduces pathology (Hultgren and Tarkowski, 2001). This is in contrast to our own results, where leptin deficiency reduces inflammation. Beyond inflammation, the effect of leptin on organ damage is also complex. While we found that leptin deficiency sustains physiological function in the face of increased organ damage during infection, thereby promoting disease tolerance, a different study showed that administration of leptin reduces remote organ injury in rats exposed to thermal burn trauma (Cakir et al., 2005). This suggests that whether leptin deficiency improves physiological function is also highly context dependent. In sum, reduction of serum leptin may serve as an important clinical tool to manage inflammation and sustain physiological function during some infections, but more work is needed to identify the specific infections that would benefit from this approach.

## AUTHOR CONTRIBUTIONS

Conceptualization: JSA

Investigation: KKS, JSA, SS, and KT.

Formal Analysis: JSA

Supervision: JSA

Writing – Original Draft: KT and JSA

Review and Editing: KT and JSA

## ACKNOWLEDGEMENTS

We thank all members of the Ayres lab for their continued support. This work was supported by a postdoctoral fellowship from the NOMIS Foundation (KT) and NIH R01 AI114929 (JSA).

## MATERIALS AND METHODS

### Mice

6-8 week old male mice were used for all experiments. C57BL/6J mice were purchased from Jackson Laboratories. *Lep^ob^* and *Lepr^db^* mice on the C57BL/6J background were purchased from Jackson Laboratories. *Il-1b^-/-^* and *Pnpla2 fabp4 cre* mice were bred in-house. All mutations were confirmed via PCR. Experiments were performed in our AAALAC-certified vivarium, with approval from The Salk Institute Animal Care and Use committee.

### Bacteria

*Yersinia pseudotuberculosis* (IP2666) was a gift from Igor Brodsky and James Bliska.

### Culturing *Yersinia pseudotuberculosis* for mouse infections

*Yersinia pseudotuberculosis* was grown overnight at 26°C on a 2xYT agar plate with 4mg/ml of irgasan (Sigma). 2xYT ingredients for 1 liter include: 16g Bacto Tryptone, 10g Bacto Yeast Extract, 5g NaCl, 15g Agar, pH to 7.0. A single colony from 2xYT/irgasan agar plate was inoculated into 6 ml of sterile 2xYT media with a 4mg/ml of irgasan. The culture was shaken overnight at 26°C (250 RPM). The next evening, a second overnight culture was started by sub-culturing the 6 ml culture in 50ml of sterile 2xYT/irgasan. 500 ml of the 6 ml culture was put into 50 ml of sterile 2xYT/irgasan media. This was shaken overnight at 26°C. The following morning an OD was taken to quantify the concentration of bacteria. Bacteria were pelleted by centrifugation and resuspended in sterile PBS to required infection concentrations. For every infection, the inoculum was serially diluted and plated in order to confirm the inocula.

### Mouse infection models

For every experiment, mice were fasted 12-16 hours overnight the day before infection. Mice were infected with 5×10^9^ CFU/ml in a 100 ml gavage. Post-infection mouse weights, food weights, and survival were assayed. Mice were single housed for all experiments.

### Food restriction

Single housed *Lep^ob^* and *Lep^db^* mice were given a restricted diet during infection that matched the feeding amounts of *Yptb*-infected WT mice each day post-infection. WT mice and control *Lep^ob^* and *Lep^db^* mice were fed *ad libitum*.

### MRI

An EchoMRI machine was used for all MRIs. MRIs were performed on uninfected and infected mice at day 1, 2 and 3 post-infection to quantify total body fat (g) and lean mass (g).

### Quantification of *Yersinia pseudotuberculosis* in mouse tissues

Spleen, mesenteric lymph nodes, cecum, colon, liver, small intestine, Peyer’s patches, were harvested from infected mice and bead beaten in PBS with 1% Triton X-100 with a BeadMill 24 benchtop bead-based homogenizer (Fisher Scientific). Tissues were serially diluted and plated on *Yersinia* selective agar containing. Agar plates were incubated at 26°C and colonies were quantified.

### ELISAs

ELISAs were performed to quantify the levels of IL-1β, IL-6, and TNFα on spleen, mesenteric lymph nodes, cecum, colon, liver, small intestine, and Peyer’s patches homogenates. All antibodies were purchased from eBioscience. For the leptin ELISA, the Mouse/Rat Leptin Quantikine ELISA Kit was used on harvested serum (R&D Systems).

### Clinical pathology

Clinical pathology analyses were performed by IDEXX Laboratories on harvested serum to measure the levels of AST, ALT, BUN, creatinine and bilirubin.

### qRT-PCR

Hypothalami were harvested and then frozen at −80°C for subsequent RNA extraction, using the PARIS™ Kit (Invitrogen). cDNA was generated using SuperScript IV Reverse Transcriptase (Invitrogen). qRT-PCR was performed using a QuantStudio 5 Real-Time PCR instrument (Applied Biosystems). Primer sequences include: *Rps17* forward 5’-CGCCATTATCCCCAGCAAG-3’ and reverse 5’-TGTCGGGATCCACCTCAATG-3’, *Lepr* forward 5’-TCATCCTACGTCTGAGCCCA-3’ and reverse 5’-GGAGTCAGGAAGGACACA CG-3’, *Fos* forward 5’-TTTATCCCCACGGTGACAGC-3’ and reverse 5’-ACACGGTCTT CACCATTCCC-3’, *Socs3* forward 5’-TTGAGCGTCAAGACCCAGTC-3’ and reverse 5’-CGTGGGTGGCAAAGAAAAGG-3’, *Npy* forward 5’-CGTGTGTTTGGGCATTCTGG-3’ and reverse 5’-AGCGGAGTAGTATCTGGCCA-3’, *Agrp* forward 5’-GCAGACCGAGCAGA AGAAGT-3’ and reverse 5’-TTGAAGAAGCGGCAGTAGCA-3’, and *Pomc* forward 5’-GTACCCCAACGTTGCTGAGA-3’ and reverse 5’-GGCTCTTCTCGGAGGTCATG-3’.

### Quantification and statistical analysis

All statistical tests were conducted using Prism version 8.0. Sample sizes, statistical tests used, and p values are indicated in each figure legend. All experiments were repeated a minimum of two times. Data shown are either a single representative experiment or data combined from independent experiments, and this is indicated in the figure legends.

---

**Supplemental Figure S1.**
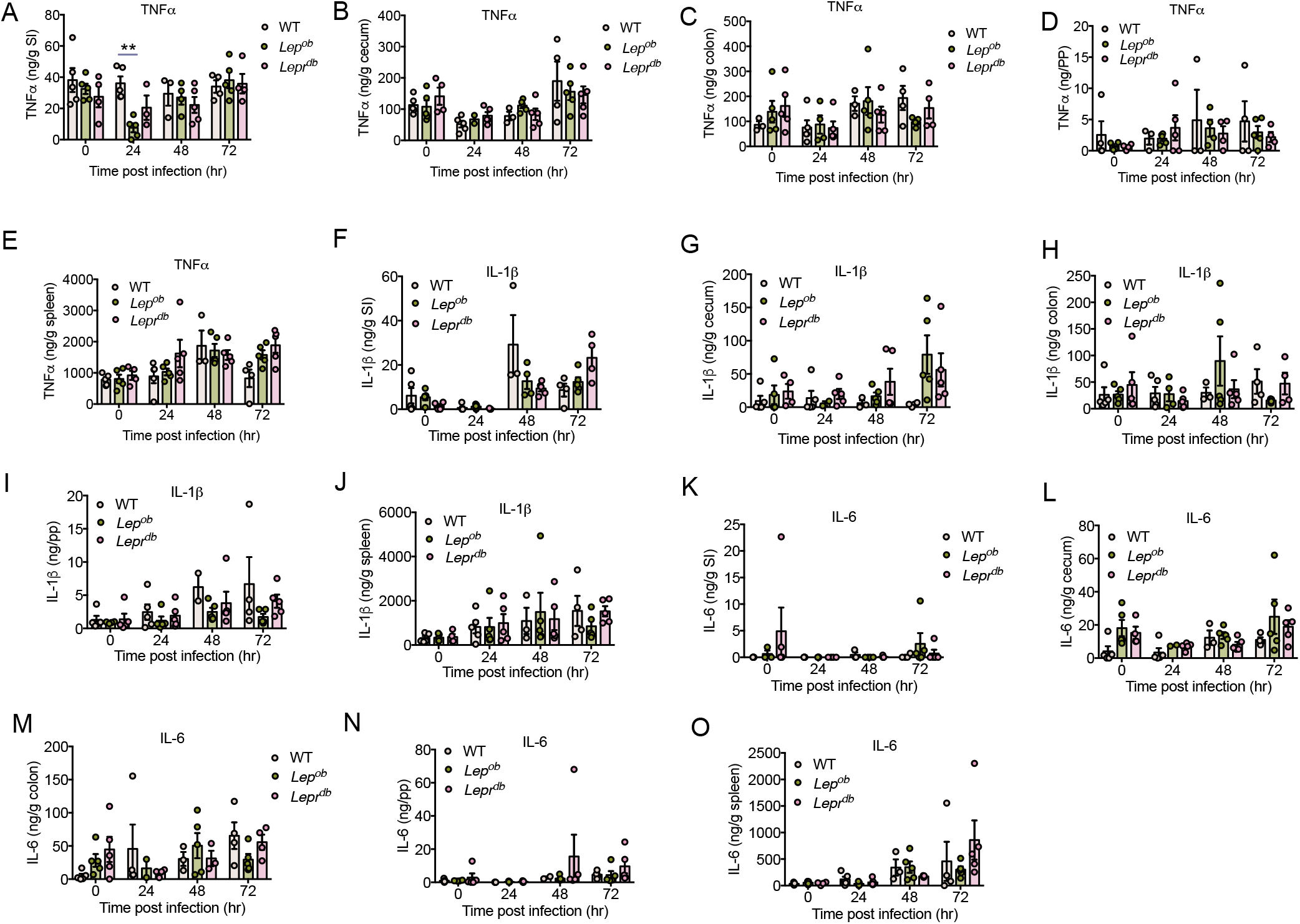
Pro-inflammatory cytokine levels in tissues during *Yptb* infection. WT, *Lep^ob^*, and *Lepr^db^* mice were infected with *Yptb* and proinflammatory cytokines in the small intestine (SI), cecum, colon, Peyer’s patches (PP) and spleen. (A) TNFα levels in SI; (B) TNFα levels in cecum; (C) TNFα levels in colon; (D) TNFα levels in PP; (E) TNFα levels in spleen; (F) IL-1β levels in SI; (G) IL-1β levels in cecum; (H) IL-1β levels in colon; (I) IL-1β levels in PP; (J) IL-1β levels in spleen; (K) IL-6 levels in SI; (L) IL-6 levels in cecum; (M) IL-6 levels in colon; (N) IL-6 levels in PP; (O) IL-6 levels in spleen. n=3-5 mice per condition. Error bars +/- SEM.

## REFERENCES

Alti, D., Sambamurthy, C., and Kalangi, S.K. (2018). Emergence of Leptin in Infection and Immunity: Scope and Challenges in Vaccines Formulation. Front Cell Infect Microbiol 8, 147.

Ayres, J.S. (2016). Cooperative Microbial Tolerance Behaviors in Host-Microbiota Mutualism. Cell 165, 1323–1331.

Ayres, J.S. (2020). The Biology of Physiological Health. Cell 181, 250–269.

Ayres, J.S., and Schneider, D.S. (2009). The role of anorexia in resistance and tolerance to infections in Drosophila. PLoS Biol 7, e1000150.

Ayres, J.S., and Schneider, D.S. (2012). Tolerance of infections. Annu Rev Immunol 30, 271–294.

Buhles, W.C., Jr., Vanderlip, J.E., Russell, S.W., and Alexander, N.L. (1981). Yersinia pseudotuberculosis infection: study of an epizootic in squirrel monkeys. J Clin Microbiol 13, 519–525.

Cakir, B., Cevik, H., Contuk, G., Ercan, F., Eksioglu-Demiralp, E., and Yegen, B.C. (2005). Leptin ameliorates burn-induced multiple organ damage and modulates postburn immune response in rats. Regul Pept 125, 135–144.

Cornelis, G.R., Boland, A., Boyd, A.P., Geuijen, C., Iriarte, M., Neyt, C., Sory, M.P., and Stainier, I. (1998). The virulence plasmid of Yersinia, an antihost genome. Microbiol Mol Biol Rev 62, 1315–1352.

Dantzer, R. (2009). Cytokine, sickness behavior, and depression. Immunol Allergy Clin North Am 29, 247–264.

Davis, K.M. (2018). All Yersinia Are Not Created Equal: Phenotypic Adaptation to Distinct Niches Within Mammalian Tissues. Front Cell Infect Microbiol 8, 261.

Dayakar, A., Chandrasekaran, S., Veronica, J., and Maurya, R. (2016). Leptin induces the phagocytosis and protective immune response in Leishmania donovani infected THP-1 cell line and human PBMCs. Exp Parasitol 160, 54–59.

Deck, C.A., Honeycutt, J.L., Cheung, E., Reynolds, H.M., and Borski, R.J. (2017). Assessing the Functional Role of Leptin in Energy Homeostasis and the Stress Response in Vertebrates. Front Endocrinol (Lausanne) 8, 63.

Dubois, A., Gervais, C., Arich, C., Gouby, A., Pignodel, C., Fabre, S., and Janbon, C. (1982). [Liver diseases associated with Yersinia infections (author’s transl)]. Nouv Presse Med 11, 1619–1621.

Farrer, W., Kloser, P., and Ketyer, S. (1988). Yersinia pseudotuberculosis sepsis presenting as multiple liver abscesses. Am J Med Sci 295, 129–132.

Fernandez-Riejos, P., Najib, S., Santos-Alvarez, J., Martin-Romero, C., Perez-Perez, A., Gonzalez-Yanes, C., and Sanchez-Margalet, V. (2010). Role of leptin in the activation of immune cells. Mediators Inflamm 2010, 568343.

Fischer, J., Gutierrez, S., Ganesan, R., Calabrese, C., Ranjan, R., Cildir, G., Hos, N.J., Rybniker, J., Wolke, M., Fries, J.W.U., et al. (2019). Leptin signaling impairs macrophage defenses against Salmonella Typhimurium. Proc Natl Acad Sci U S A 116, 16551–16560.

Francisco, V., Pino, J., Campos-Cabaleiro, V., Ruiz-Fernandez, C., Mera, A., Gonzalez-Gay, M.A., Gomez, R., and Gualillo, O. (2018). Obesity, Fat Mass and Immune System: Role for Leptin. Front Physiol 9, 640.

Galindo, C.L., Rosenzweig, J.A., Kirtley, M.L., and Chopra, A.K. (2011). Pathogenesis of Y. enterocolitica and Y. pseudotuberculosis in Human Yersiniosis. J Pathog 2011, 182051.

Grunfeld, C., Zhao, C., Fuller, J., Pollack, A., Moser, A., Friedman, J., and Feingold, K.R. (1996). Endotoxin and cytokines induce expression of leptin, the ob gene product, in hamsters. J Clin Invest 97, 2152–2157.

Hart, B.L. (1988). Biological basis of the behavior of sick animals. Neuroscience & Biobehavioral Reviews 12, 123–137.

Heroven, A.K., and Dersch, P. (2014). Coregulation of host-adapted metabolism and virulence by pathogenic yersiniae. Front Cell Infect Microbiol 4, 146.

Hsu, A., Aronoff, D.M., Phipps, J., Goel, D., and Mancuso, P. (2007). Leptin improves pulmonary bacterial clearance and survival in ob/ob mice during pneumococcal pneumonia. Clin Exp Immunol 150, 332–339.

Hultgren, O.H., and Tarkowski, A. (2001). Leptin in septic arthritis: decreased levels during infection and amelioration of disease activity upon its administration. Arthritis Res 3, 389–394.

Kim, Y.K., Cho, M.H., Hyun, H.S., Park, E., Ha, I.S., Cheong, H.I., and Kang, H.G. (2019). Acute kidney injury associated with Yersinia pseudotuberculosis infection: Forgotten but not gone. Kidney Res Clin Pract 38, 347–355.

Luan, H.H., Wang, A., Hilliard, B.K., Carvalho, F., Rosen, C.E., Ahasic, A.M., Herzog, E.L., Kang, I., Pisani, M.A., Yu, S., et al. (2019). GDF15 Is an Inflammation-Induced Central Mediator of Tissue Tolerance. Cell 178, 1231–1244 e1211.

Mancuso, P., Gottschalk, A., Phare, S.M., Peters-Golden, M., Lukacs, N.W., and Huffnagle, G.B. (2002). Leptin-deficient mice exhibit impaired host defense in Gram-negative pneumonia. J Immunol 168, 4018–4024.

Mastronardi, C.A., Yu, W.H., Srivastava, V.K., Dees, W.L., and McCann, S.M. (2001). Lipopolysaccharide-induced leptin release is neurally controlled. Proc Natl Acad Sci U S A 98, 14720–14725.

Mikula, K.M., Kolodziejczyk, R., and Goldman, A. (2012). Yersinia infection tools-characterization of structure and function of adhesins. Front Cell Infect Microbiol 2, 169.

Mohammadi, S., and Isberg, R.R. (2009). Yersinia pseudotuberculosis virulence determinants invasin, YopE, and YopT modulate RhoG activity and localization. Infect Immun 77, 4771–4782.

Moore, S.I., Huffnagle, G.B., Chen, G.H., White, E.S., and Mancuso, P. (2003). Leptin modulates neutrophil phagocytosis of Klebsiella pneumoniae. Infect Immun 71, 4182–4185.

Otero, M., Lago, R., Gomez, R., Dieguez, C., Lago, F., Gomez-Reino, J., and Gualillo, O. (2006). Towards a pro-inflammatory and immunomodulatory emerging role of leptin. Rheumatology (Oxford) 45, 944–950.

Pan, W.W., and Myers, M.G., Jr. (2018). Leptin and the maintenance of elevated body weight. Nat Rev Neurosci 19, 95–105.

Prasad, N., and Patel, M.R. (2018). Infection-Induced Kidney Diseases. Front Med (Lausanne) 5, 327.

Raberg, L., Graham, A.L., and Read, A.F. (2009). Decomposing health: tolerance and resistance to parasites in animals. Philos Trans R Soc Lond B Biol Sci 364, 37–49.

Radigan, K.A., Morales-Nebreda, L., Soberanes, S., Nicholson, T., Nigdelioglu, R., Cho, T., Chi, M., Hamanaka, R.B., Misharin, A.V., Perlman, H., et al. (2014). Impaired clearance of influenza A virus in obese, leptin receptor deficient mice is independent of leptin signaling in the lung epithelium and macrophages. PLoS One 9, e108138.

Rao, S., Schieber, A.M., O’Connor, C.P., Leblanc, M., Michel, D., and Ayres, J.S. (2017). Pathogen-Mediated Inhibition of Anorexia Promotes Host Survival and Transmission. Cell 168, 503–516 e512.

Sanchez, K.K., Chen, G.Y., Schieber, A.M.P., Redford, S.E., Shokhirev, M.N., Leblanc, M., Lee, Y.M., and Ayres, J.S. (2018). Cooperative Metabolic Adaptations in the Host Can Favor Asymptomatic Infection and Select for Attenuated Virulence in an Enteric Pathogen. Cell 175, 146–158.e115.

Schieber, A.M.P., Lee, Y.M., Chang, M.W., Leblanc, M., Collins, B., Downes, M., Evans, R.M., and Ayres, J.S. (2015). Disease tolerance mediated by microbiome E. coli involves inflammasome and IGF-1 signaling. Science (New York, NY) 350, 558–563.

Schneider, D.S., and Ayres, J.S. (2008). Two ways to survive infection: what resistance and tolerance can teach us about treating infectious diseases. Nat Rev Immunol 8, 889–895.

Schwartz, M.W., Seeley, R.J., Woods, S.C., Weigle, D.S., Campfield, L.A., Burn, P., and Baskin, D.G. (1997). Leptin increases hypothalamic pro-opiomelanocortin mRNA expression in the rostral arcuate nucleus. Diabetes 46, 2119–2123.

Souza-Almeida, G., D’Avila, H., Almeida, P.E., Luna-Gomes, T., Liechocki, S., Walzog, B., Hepper, I., Castro-Faria-Neto, H.C., Bozza, P.T., Bandeira-Melo, C., et al. (2018). Leptin Mediates In Vivo Neutrophil Migration: Involvement of Tumor Necrosis Factor-Alpha and CXCL1. Front Immunol 9, 111.

Sucajtys-Szulc, E., Turyn, J., Goyke, E., Korczynska, J., Stelmanska, E., Slominska, E., Smolenski, R.T., Rutkowski, B., and Swierczynski, J. (2010). Differential effect of prolonged food restriction and fasting on hypothalamic malonyl-CoA concentration and expression of orexigenic and anorexigenic neuropeptides genes in rats. Neuropeptides 44, 17–23.

Tian, Z., Sun, R., Wei, H., and Gao, B. (2002). Impaired natural killer (NK) cell activity in leptin receptor deficient mice: leptin as a critical regulator in NK cell development and activation. Biochem Biophys Res Commun 298, 297–302.

Trites, M.J., and Clugston, R.D. (2019). The role of adipose triglyceride lipase in lipid and glucose homeostasis: lessons from transgenic mice. Lipids Health Dis 18, 204.

Troha, K., and Ayres, J.S. (2020). Metabolic Adaptations to Infections at the Organismal Level. Trends Immunol 41, 113–125.

Tschop, J., Nogueiras, R., Haas-Lockie, S., Kasten, K.R., Castaneda, T.R., Huber, N., Guanciale, K., Perez-Tilve, D., Habegger, K., Ottaway, N., et al. (2010). CNS leptin action modulates immune response and survival in sepsis. J Neurosci 30, 6036–6047.

Turner, S.M., Roy, S., Sul, H.S., Neese, R.A., Murphy, E.J., Samandi, W., Roohk, D.J., and Hellerstein, M.K. (2007). Dissociation between adipose tissue fluxes and lipogenic gene expression in ob/ob mice. Am J Physiol Endocrinol Metab 292, E1101–1109.

Viboud, G.I., and Bliska, J.B. (2005). Yersinia outer proteins: role in modulation of host cell signaling responses and pathogenesis. Annu Rev Microbiol 59, 69–89.

Wang, A., Huen, S.C., Luan, H.H., Yu, S., Zhang, C., Gallezot, J.D., Booth, C.J., and Medzhitov, R. (2016). Opposing Effects of Fasting Metabolism on Tissue Tolerance in Bacterial and Viral Inflammation. Cell 166, 1512–1525 e1512.

Wang, B., Chandrasekera, P.C., and Pippin, J.J. (2014a). Leptin- and leptin receptor-deficient rodent models: relevance for human type 2 diabetes. Curr Diabetes Rev 10, 131–145.

Wang, X., Parashar, K., Sitaram, A., and Bliska, J.B. (2014b). The GAP activity of type III effector YopE triggers killing of Yersinia in macrophages. PLoS Pathog 10, e1004346.

